# Sequential chromogenic immunohistochemistry: spatial analysis of lymph nodes identifies contact interactions between plasmacytoid dendritic cells and plasmablasts

**DOI:** 10.1101/2023.04.13.536793

**Authors:** N.M. Claudio, M.T. Nguyen, A.A. Wanner, F. Pucci

**Author notes:** Conflict of interest statement: AAW is founder and share-holder of ariadne.ai AG.

## Abstract

Recent clinical observations highlight the importance of the spatial organization of immune cells into lymphoid structures for the success of cancer immunotherapy and patient survival. Sequential chromogenic immunohistochemistry (scIHC) supports the analysis of multiple biomarkers on a single tissue section thus providing unique information about relative location of cell types and assessment of disease states. Unfortunately, widespread implementation of scIHC is limited by lack of a standardized, rigorous guide to the development of customized biomarker panels and by the need for user-friendly analysis pipelines able to streamline the extraction of meaningful data. Here, we examine major steps from classical IHC protocols and highlight the impact they have on the scIHC procedure. We report practical examples and illustrations of the most common complications that can arise during the setup of a new biomarker panel and how to avoid them. We described in detail how to prevent and detect cross-reactivity between secondary reagents and carry over between detection antibodies. We developed a novel analysis pipeline based on non-rigid tissue deformation correction, Cellpose-inspired automated cell segmentation and computational network masking of low-quality data. The resulting biomarker panel and pipeline was used to study regional lymph nodes from head and neck cancer patients. We identified contact interactions between plasmablasts and plasmacytoid dendritic cells *in vivo*. Given that TLR receptors, which are highly expressed in plasmacytoid dendritic cells play a key role in vaccine efficacy, the significance of this cell-cell interaction decisively warrants further studies. In conclusion, this work streamlines the development of novel biomarker panels for scIHC, which will ultimately improve our understanding of immune responses in cancer.

## 1. Introduction

The clinical success of immunotherapeutic approaches for cancer patients warrants a better understanding of immune responses in cancer, in order to increase patient outcomes and decrease adverse reactions. Biomarker discovery and assessment is a key tool in modern immune-oncology practice [1]. Defining which biomarkers are useful as prognostic and predictive indicators of disease course and therapeutic outcome is not only intrinsically challenging due to the complexity of the immune system, but also because no single marker has been validated to correlate with clinical response [2, 3]. The need to quantify multiple biomarkers is especially important in the context of immune responses to cancer, which summons both adaptive and innate immune cells to both the tumor microenvironment and sentinel lymph nodes [4]. Recent advances in our understanding of the interplay between cancer and the immune system highlight the importance of the spatial organization of immune cells, adding another layer of complexity to the problem [5–9]. Thus, innovative wet and dry lab techniques to tackle these challenges are urgently needed.

Sequential chromogenic immunohistochemistry (scIHC) is a novel experimental and analytical approach that allows researchers to assess multiple (>10) biomarkers on a single tissue section. This technique promises to vertically innovate biomarker discovery and assessment of disease states [10]. Contrary to multiplex IHC, in which two or more biomarkers are detected at the same time, biomarkers in a sequential IHC panel are tested one at a time [11]. Hence, the cells present on the slide can be sampled for tens of biomarkers, such as it is common practice in flow cytometry, with the added advantage of retaining the relative location to each other [10]. Once data is collected, the co-expression of various markers within the same cell and its spatial relations with nearby immune and tumor cells can be quantified [13]. These features grant scIHC a spot in the family of approaches commonly referred to as “tissue cytometry” [12]. Compared to other tissue cytometry approaches, the technique described here is not limited to 5-10 biomarkers (such as MICSSS and SIMPLE, reviewed in [12]). Fluorescence-based protocols (such as cIF and MxIF) can be somewhat less time-intensive than scIHC, but are not integrated in pathology labs and some methods (like CODEX and NanoString) require special devices and reagents [12]. The consumables employed during scIHC are commercially available and widely used in clinical pathology laboratories, which makes scIHC a simple and affordable technique as compared to other equivalent approaches [14]. To benefit patients, the transition into the clinic will require the reduction of large, research-grade biomarker panels down to the minimum number of biomarkers strictly necessary for clinical decision making.

Our group recently employed scIHC to quantify the tumor immune microenvironment of sinonasal squamous cell carcinoma (SNSCC). We identified potential prognostic biomarkers in a cohort of 38 patients by showing increased T-cell populations and decreased myeloid-cell populations in SNSCC patients without recurrent disease, as compared with recurring patients [15]. Although the characterization of the tumor immune microenvironment can inform treatment decisions and provide useful prognosticators [5, 16–19], the unpredictable nature of the tumor microenvironment can represent a significant obstacle to reproducibly and accurately assessing immune responses [20–22]. Moreover, polyclonal antibodies cannot generally be employed to assess the tumor immune microenvironment, likely due to the presence of aberrant cells carrying different glycosylations and oxidized epitopes that greatly increase background signal (unpublished observations). On the other hand, regional lymph nodes are highly structured organs that are inhabited by normal immune cells responsible for key immune reactions to tumor antigens [23]. In addition, immune cell location within highly compartmentalized lymph nodes can be indicative of function [24, 25]. Lastly, sentinel node biopsy and lymph node dissection are standard of care in patients with early disease (T1-2) and with evident nodal disease at the time of diagnosis, respectively [26–28]. For these reasons, analysis of patient lymph nodes represents a promising alternative to inform treatment decisions and provide important prognostic indications.

To develop a novel scIHC panel, it is not enough to simply add single IHC steps back-to-back. Several considerations into reagent selection and compatibility need to be assessed in order to avoid getting into a daunting task riddled with unexpected complications. Here, we examine major steps from classical IHC protocols and highlight the impact they have on the scIHC procedure. As a case study, we describe the development of a novel scIHC panel, with special focus on staining order, on how to avoid signal carryover and secondary reagent cross-reactivity, and on comparison between alternative procedures. Findings for each antibody tested are provided in an external resource to guide development of other customized panels. We use this scIHC panel to quantify immune parameters in regional lymph nodes from patients diagnosed with head and neck cancer (HNC; Table 1), that we analyzed by developing a novel bio- informatic pipeline. Together, our findings provide a systematic and practical approach to develop novel biomarker panels for sequential histological studies that will contribute to improve our understanding of immune responses in cancer patients.

**Table 1:** Patient characteristics (n=10)

| Characteristic: | N(%)<br>Overall |
| --- | --- |
| Age at diagnosis | 58.6 (14.15)* |
| Female | 3 |
| Male | 7 |
| Race: White/Caucasian | 10 |
| Race: African American | 0 |
| **Overall Anatomic Stage at dx:** |  |
| Stage-1 | 0 |
| Stage-2 | 1 |
| Stage-3 | 1 |
| Stage-4 | 8 |
| **Tumor location:** |  |
| Neck | 3 |
| Tongue/Maxilla | 4 |
| Larynx/Pharynx/Tonsil | 3 |
| **Nodal status** |  |
| 0 (no nodal metastasis) | 1 |
| 1 (metastases in 1-3 nodes) | 3 |
| 2 (metastases in ≥4 nodes) | 6 |
| **Treatment type:** |  |
| Surgery alone | 3 |
| Surgery + radiation | 5 |
| Surgery + CRT | 2 |

\*Average (SD)

## 2. Materials and methods

### 2.1 Clinical Samples

Formalin-fixed paraffin embedded (FFPE) tissue sections (5µm) of tumor-free N0 regional lymph nodes from anonymized patients diagnosed with HPV+ or HPV– squamous cell carcinoma of the head and neck were obtained from the Oregon Health and Science University (OHSU) Knight Cancer Institute Biolibrary (IRB #19903).

### 2.2 Sequential chromogenic Immunohistochemistry and image acquisition

Our protocol was guided by the seminal work of Tsujikawa et. al. [11, 29] with some modifications. FFPE patient tissue sections were baked for 60°C for 60 minutes total, with a 5-minute incubation at room temperature after the first 30 minutes. These slides were then deparaffinized by submerging them in xylene for 5 minutes, and then repeated again for another 5 minutes. Slides were then rehydrated by incubating in serially graded ethanol, with a final incubation in distilled water. An initial counterstain of Hematoxylin (Dako, S3301) was performed for 1 minute, with washes in distilled water, and placed in Tris Buffered Saline with 2% Tween, pH 7.4 (Boston BioProducts, IBB-180X). Slides were coverslipped (Corning, 2975246) and imaged using an Aperio ImageScope AT (Leica Biosystems) at 20X magnification. Coverslips were removed by gentle agitation of the slides in Tris Buffered Saline with 2% Tween (TBST). Slides were then subjected to heat-mediated antigen retrieval by placing slides in boiling Citrate buffer with a pH of 6.0 (Abcam, ab93678) and steamed for 30 minutes (100°C). Slides were allowed to cool to room temperature, washed with distilled water, and placed in TBST. Endogenous peroxidase activity was blocked by incubating slides with Dako dual endogenous peroxidase block (Dako, S2003) for 10 minutes at room temperate. Slides were washed with distilled water, placed in TBST for 1 minutes. Additional protein blocking was performed by incubating slides in 1X Phosphate buffered saline (PBS) (Corning, 21- 040-CM) containing 5% horse serum and 2.5% Bovine Serum Albumin for 10 minutes at room temperature. Slides experienced primary antibodies at saturating dilutions, with incubation times and temperatures previously optimized during testing (Table S2). After primary antibody incubation, slides were washed in TBST, and incubated with an appropriate F(ab’) fragment–specific secondary-antibody– labeled polymer conjugated to horseradish peroxidase for 30 minutes at room temperature (Table S3). After primary and secondary antibody detections steps, signal was visualized using an alcohol-soluble peroxidase substrate 3-amino-9-ethylcarbazole (AEC) (Vector Labs, SK-4200), followed by whole-slide digital scanning. After imaging, AEC was removed using graded ethanol incubations, briefly washed in distilled water, and placed in TBST. Secondary HRP signal was either inactivated by performing two blocking steps of Dako dual endogenous peroxidase block (Dako, S2002) for 10 minutes at room temperature (each), in addition to a protein blocking step to allow for another primary antibody produced in a different species to be applied as a different round, or antibodies were stripped in heated citrate buffer (pH 6.0) to begin a new cycle.

In the last cycle of the scIHC panel, a Tris-EDTA antigen retrieval buffer, pH 9.0 (Abcam, ab93684) was required for efficient signal detection. Antibodies from the previous cycle required stripping in a Citrate buffer, as mentioned above, before “conditioning” the tissue with the EDTA based buffer. This “conditioning” was also heat-mediated, and steamed for 30 minutes. The protocol for staining and visualization was performed as in previous cycles, including both primary and secondary antibodies. Antibodies that require a Tris-EDTA antigen retrieval can also be placed in rounds at the first cycle(s) if successive heat treatment adversely affects the tissue integrity on the slide. Changing antigen retrieval buffers within and experiment should be avoided; however, this method can be utilized if that is not possible, as was the case for this experiment. At the end of the last cycle, AEC was removed from the slides using ethanol gradient washes with, a final counterstain with Hematoxylin was performed, and slides were imaged.

### 2.3 Database for optimization of antibody testing

In addition to determining an antibody’s saturating dilution and incubation parameters, it is necessary to identify its appropriate location within the panel. Ideally, an antibody of interest would be tested in each round of every cycle, however due to budgetary and temporal constraints, this may not be possible. To overcome such constraints, we created and utilized a detailed database (AirTable Inc., San Francisco CA) to track all tests in an effort to optimize the development of the scIHC panel (Table S4). As tests and experimental data are generated, the database is continually updated to streamline the creation of future scIHC panels, and currently has >300 entries. The database can be accessed at [available upon acceptance of the manuscript].

All slides were kept in TBST with 0.02% sodium azide and used as a repository of tissue to test at different cycles. A separate database was used to track slides, their tissue type, their, in addition to the cycle and round number. If an antibody was tested and determined to work at cycle 1 and again at cycle 5, the authors assumed this antibody would also work in cycles 2-4. Moreover, if a specific antibody did not work at a given cycle, it was assumed it would also not work in subsequent cycles.

### 2.4 Image processing

For efficient, parallel handling and processing of the large scIHC images, the data was first’ chunked into a multi-resolution, multi-channel format of small image squares of 512 x 512 pixels, compatible with the open-source multi-channel volume image annotation tool KNOSSOS (https://knossos.app) [30]. KNOSSOS has originally been developed for serial section 3D electron microscopy data, but it can also be used with bidimensional data, which is a 3D image with just one slice. Subsequently, each IHC image was registered to the HEM image. Alignment between all the biomarker images for each section is necessary because the scIHC procedure causes shifts and micro-deformations in the tissue. Our computational methods correct for both local distortions and macro-shifts of the tissues on the slide (Figure S2). After a global rigid alignment at downsampled resolution, local non-linear distortions were corrected chunk-wise as follows. First, correspondence point candidates were automatically determined in the IHC images and the HEM image using a customized normalized cross-correlation procedure [31]. Next, correspondence point candidates with locally non-consistent distortion directions and amplitudes were pruned and a distortion field was fitted to the remaining set of correspondence points using fast natural neighbor interpolation. An example of the alignment outcome is provided in Figure S3. In order to ignore low quality data, including out of focus or damaged areas, as well as cells from nearby tissues that may express some of the biomarkers used but that do not belong to the analysis (eg. CD141 is also expressed in endothelial cells in fat tissue), we trained the computational network to detect and mask artifacts like folds, out-of-focus areas, red blood cells within vessels and non-lymph node tissue (Figure S4). Cell outlines were segmented in the hematoxylin image (before cycle 1) using a parallelized Cellpose-based segmentation workflow [32]. To maximize the segmentation accuracy, the network was iteratively re-trained with targeted human generated ground truth. Cell segmentation accurately detected most nuclei irrespective of their shapes, including round lymphocyte nuclei and elongated macrophage nuclei (Movie S1). Regions of interest have been determined automatically using a tailored U-net based artifact and tissue segmentation workflow. Within the regions of interest, each segmented cell was assigned a unique ID and the average intensity of the magenta color channel within the segmented cell was calculated for all registered IHC images. The final output of the pipeline was a list of unique cell IDs with x/y coordinates and average intensities for each of the scIHC markers, stored in FCS format for import in Cytobank (community version, BD), a web-based flow cytometry data analysis application. Subsets of interest (see below) were gated in Cytobank based on biomarker intensities. Each gated cell subset was then exported as a list of cells defined by x/y coordinates. These lists were visualized in KNOSSOS for visual inspection of the results (cell segmentation and biomarker presence). For spatial analysis, each cell within the plasmablast subset (defined by CD138 expression) was assigned the median Euclidean distance to the nearest 5 cells from the following subsets: plasmacytoid dendritic cells (pDCs: BDCA2), type-1 and type-2 conventional dendritic cells (cDC1: CD141+CD1c–; cDC2: CD141–CD1c+), sinusoidal macrophages (CD169), non-sinusoidal macrophages (CD169–CD68+), NK cells (CD56) and granulocytes (CD66b). Frequency distributions of nearest neighbors was plotted with Graphpad Prism (version 9).

## 3. Results

An overview of the scIHC protocol is depicted in Figure 1, along with the indication of which steps of the protocol were improved, as described below.

**Figure 1.**
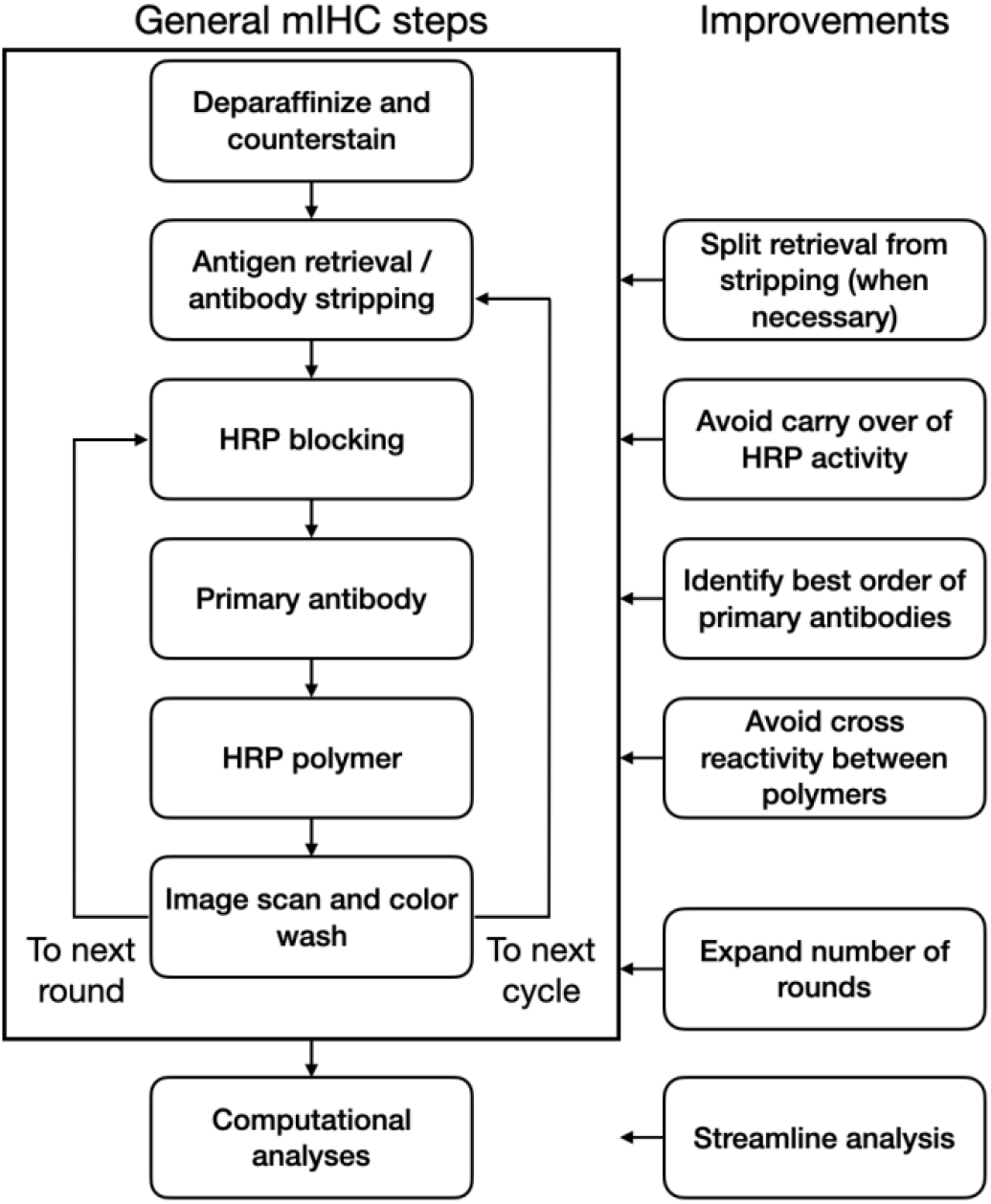
Schematic of the mIHC process and the improvements developed in this work.

### 3.1 Determining staining order in a panel

Once a list of biomarkers of interest is defined, the first step in designing a scIHC panel is testing the individual antibodies against those biomarkers. The number of cycles needed is determined by the highest number of primary antibodies from the same species: for example, if a list of 20 primary antibodies contains 8 made in mouse, 10 in rabbit and 2 in rat, 10 cycles will be needed. In order to simplify development of new panels and avoid testing of primary antibodies at each stripping cycle, we collected all our testing into a detailed database for other investigators to use (see methods). The need to identify the latest cycle at which the primary antibodies still work comes from the fact that multiple stripping cycles likely affect epitope conformation and subsequent ability of the primary antibody to detect its target. Some antigens are preserved across many cycles (see below), while others are more sensitive to the relatively harsh condition necessary for stripping primary antibodies from the previous cycle. Luckily, multiple antibodies can be assigned to the same order position (that is, the same stripping cycle) if they can be placed in different rounds within that stripping cycle (that is, they are from different species), which greatly simplifies primary antibody order assignment. Antigens with epitopes sensitive to stripping should be prioritized in earlier cycles [14]. Thus, determining the staining order is crucial to identify each primary antibody’s order within the panel. To this end, practice slides should be reused as the various primary antibodies are tested to identify ideal staining conditions, including incubation time/temperature and dilution factor. This way, information on staining efficiency in several different cycles can be obtained. For example, a mouse anti-human CD66b (see Table S1) was determined to work at a dilution of 1:600 for 30 minutes at room temperature at cycle 1 (Figure 2A). The slide was stripped and used to identify ideal staining conditions for a rabbit anti-human CD141, and subsequently stripped again for assessment of other primary antibodies. The same mouse anti-human CD66b was then retested at cycle 9 under the identified optimal conditions (Table S1) but no staining was observable (Figure 2B). Similarly, the rabbit anti-human CD141 was determined to work at a dilution of 1:100 overnight at 4°C in cycle 2 (Figure 2C), but not at cycle 9 (Figure 2D). Stripping resistance of a biomarker can be defined as the ability of one of its epitopes to restore its conformation for binding to a given antibody. Biomarkers resistant to multiple stripping cycles can be found. For example, a mouse anti-human CD68 and a rabbit anti-human BCL6 antibodies worked well in cycle 11 (data not shown). All data from these antibody tests was collected into a detailed database (see methods), in a searchable format, that allows to determine a tentative order in which antibodies could be placed into a new panel, without the need to systematically assess each antibody at every cycle. Collecting this information in a database is useful for determining cycle placement when developing new panels, such that testing could be tailored based on the data already available, thereby greatly reducing the amount of work necessary.

**Figure 2.**
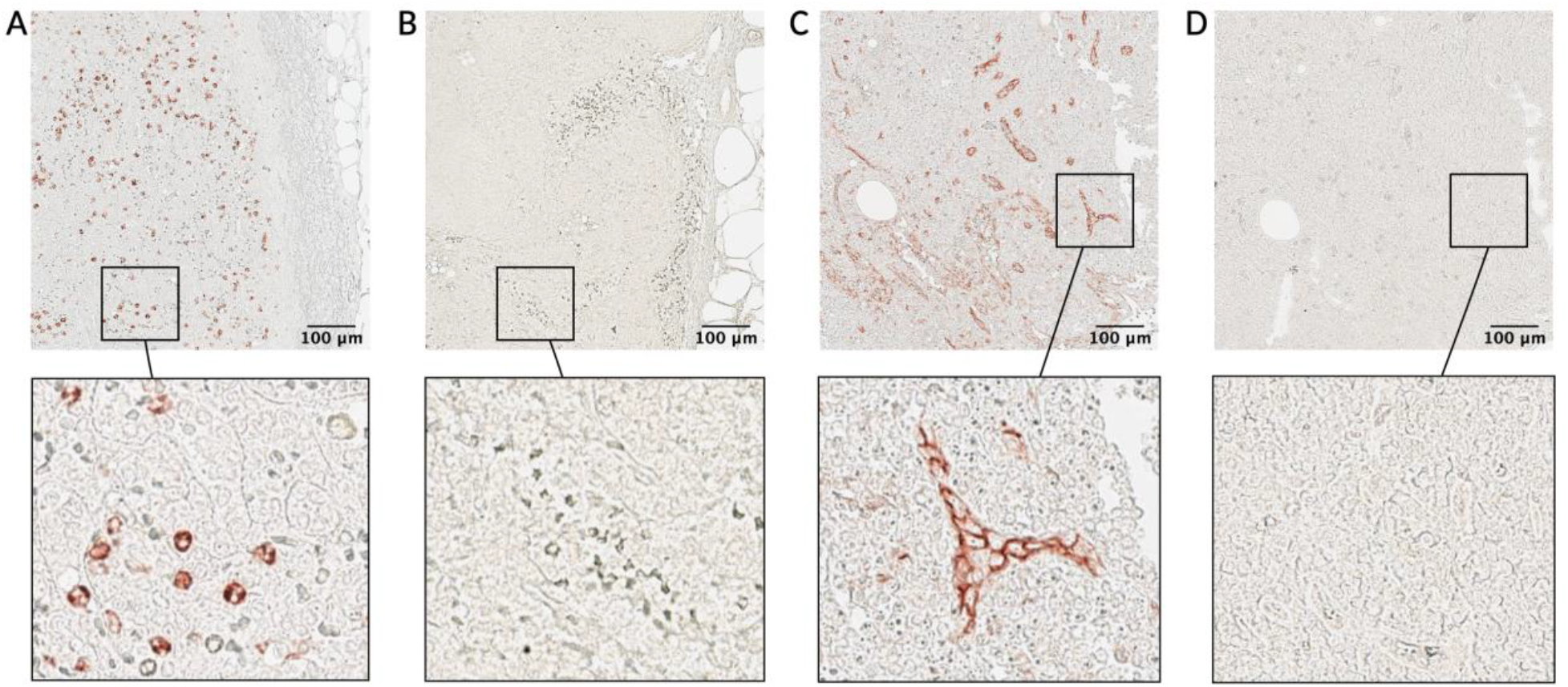
Cycle-dependent performance of antibodies. Human lymph node histological slides were stained with CD66b at a dilution of 1:600 for 30 minutes at room temperature at cycle 1 (A) and cycle 9 (B). Additionally human lymph node samples were stained with CD141 at a dilution of 1:100 overnight at 4°C at cycle 2 (C) and at cycle 9 (D). After testing, it was determined that these antibodies cannot be used at later cycles within these specific staining conditions. Sub-capsular region is shown in both (A) and (B). Consecutive sections are shown in (C) and (D). Top row scale bar: 100 um (5X magnification).

### 3.2 Expanded selection of secondary reagents

In addition to optimization of staining conditions for primary antibodies to be used in a scIHC panel, it is also necessary to have reliable secondary reagents for efficient biomarker detection. In order to maximize the number of biomarkers within each stripping cycle (see above), we expanded the repertoire of anti-human primary antibodies to include less common species, such as goat, rat and sheep (mouse and rabbit anti-human primary antibodies arguably are the most common). These primary antibodies are used sequentially within a cycle. To match these additional primary antibodies with their corresponding secondary reagents, we compared traditional HRP-conjugated antibodies (IgG) with HRP polymers (Fab’). Although polymeric secondary antibodies are widely used in IHC, a formal comparison with their monomeric counterparts in scIHC has not been assessed. These tests are especially important given the increased potential for cross-reactivity of secondary polymeric reagents used sequentially, as we discuss in Section 3.4. Even after optimizing dilutions and incubation times, we found that detection using an HRP-conjugated IgG showed dimmer staining as compared to an HRP polymer, independent of biomarker identity and while maintaining the same primary antibody conditions (Figure 3). The secondary HRP polymers not only showed strong staining in early cycles (data not shown), they also demonstrated better signaling in later cycles. For example, CD11c detection using the traditional secondary HRP-conjugated IgG at cycle 1 (Figure 3A) was less intense than a secondary HRP polymer using the same primary antibody conditions at cycle 4 (Figure 3B). The difference was more pronounced when observing Tbet signaling, with the secondary HRP-conjugated IgG showing dim stain at cycle 3 (Figure 3C), but the secondary HRP polymer showing a much stronger signal at cycle 11 (Figure 3D). In conclusion, HRP polymers outperform traditional HRP-conjugated IgG not only because they are ready to use and save time, but also because these reagents provide increased signal intensity and reduced background.

**Figure 3.**
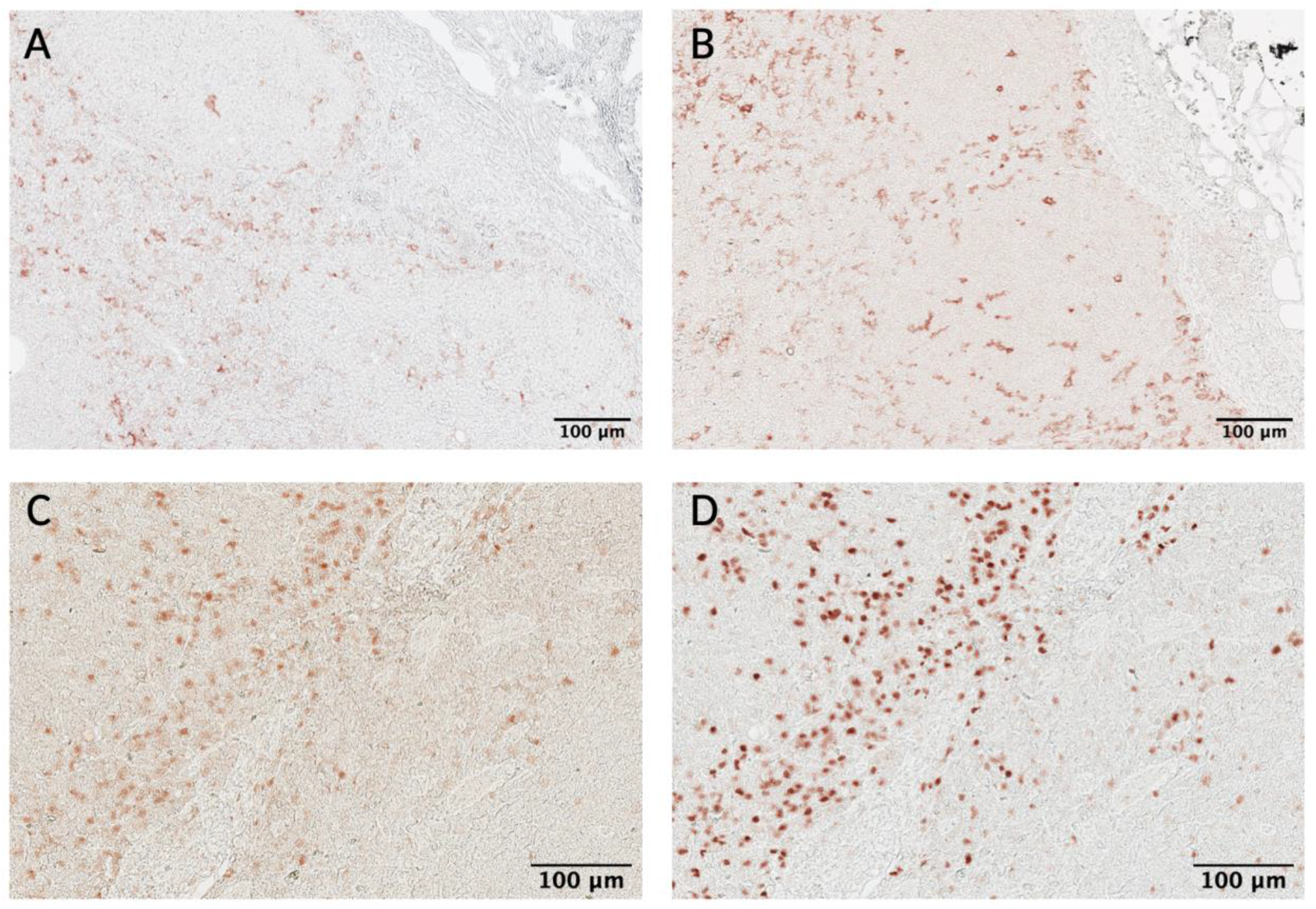
Polymeric secondary reagents outperformed traditional secondary antibodies. Primary antibodies used were rabbit anti-human CD11c (A, B) and rabbit anti-human Tbet (C, D). Detection using a donkey anti-rabbit HRP antibody (A, C) showed dimmer staining as compared to detection using a horse (B) or goat (D) anti-rabbit HRP polymer. Sub-capsular region is shown in both (A) and (B). Same section is shown in (C) and (D). Scale bar: 100 um (10X).

### 3.3 Avoiding carryover within a stripping cycle

Once each primary antibody has been assigned to a stripping cycle, preliminary testing of the whole scIHC panel can start. During this final testing phase, special consideration should be paid to signal carryover within each stripping cycle. We define carryover as signal from the previous round appearing in the current round. This can happen because primary antibodies and secondary reagents are consecutively added to the section during each round, without stripping (Figure 1). Thus, secondary reagents may form chromogenic signal where previously used primary antibodies are bound. As a result, the antibody signal cannot be trusted because it would be a mix of multiple markers. It is important to eliminate carryover in scIHC to ensure accurate data acquisition and analysis. To detect carryover, we routinely checked for similarities in signal patterns between the current and the previous rounds. During testing of patient samples, we had an interesting case of selective carryover in some patient samples but not others which allowed us to investigate the issue in detail (Figure 4). More specifically, a goat anti-human CD20 was used in the second round. Appropriate signal was detected in all patient samples (See Figure 4A and 4E). A mouse anti-human CD66b antibody was used in the third round. Most patient samples showed identical CD20-like signal as the previous round that we interpreted as carryover (Figure 4B); however, one patient had the expected CD66b staining (Figure 4F). This created an opportunity to test conditions that eliminate carry over, and recover accurate signal for data acquisition. We reasoned that carryover is due to insufficient blocking of peroxidase from the previous round, which involved a very abundant marker, CD20. To determine if extra peroxidase blocking would aid in avoiding signal carry over of goat anti-human CD20 into the round with mouse anti-human CD66b, AEC from all patient slides was removed and endogenous peroxidase activity was blocked twice. To confirm absence of peroxidase activity, we applied AEC to the slides immediately after the double blocking step above and imaged the slides. After confirming complete absence of peroxidase activity (Figure 4C and 4G), we proceeded with staining for CD66b and its secondary reagent. Upon AEC visualization, we observed the expected signal –albeit somewhat dimmer, in the patient sample that did not have carryover (Figure 4H), suggesting that double peroxidase treatment does not prevent detection of biomarkers. Importantly, we observed a different staining pattern that corresponded to CD66b signaling in patient samples that previously contained carryover (Figure 4D), suggesting that, at least for abundant markers like CD20 in lymph nodes, double peroxidase blocking is an effective strategy to prevent carry over.

**Figure 4.**
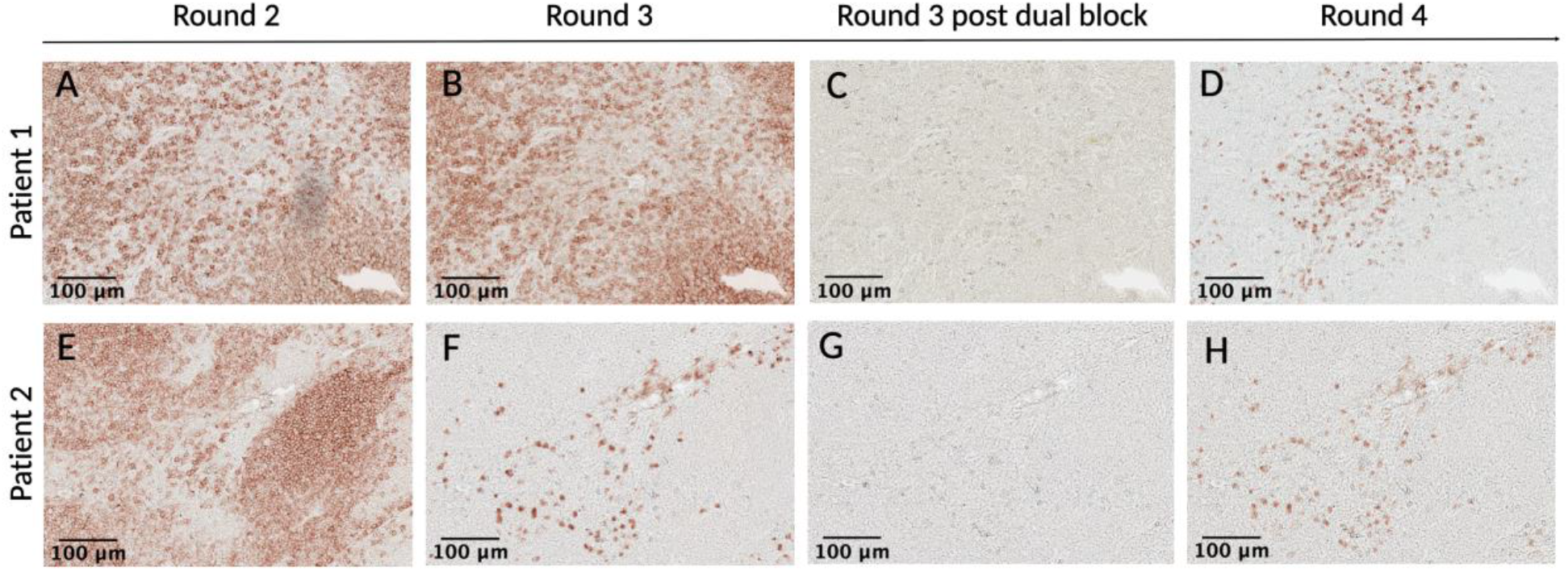
Two-step peroxidase blocking avoids signal carry over. Patient samples were stained with CD20 in cycle 1 round 2 (A and E), and subsequently stained with CD66b in cycle 1 round 3 (B and F). Appropriate signal was detected in one patient sample (F), but carry over was detected in the other (B). The AEC substrate was removed, and samples were blocked twice with endogenous peroxidase block, and developed again with AEC to ensure complete removal of signal (C and G). Slides were then re-stained with CD66b, and the appropriate secondary HRP polymer. Upon AEC visualization, CD66b signal was recovered (D) and the same CD66b staining pattern was observed in the sample with no carry over (H). Same section is shown in (A-D) and in (E-H). Scale bar: 100 um (10X).

### 3.4 Avoiding secondary reagent cross-reactivity within a stripping cycle

The use of primary antibodies from different species allows greater flexibility in assigning an order to each of them in the panel by placing multiple biomarkers in the same stripping cycle. Due to the variety of species we used in each cycle, we were primarily concerned with cross-reactivity between secondary reagents. Although methods to avoid cross-reactivity between secondary antibodies in multicolor IHC approaches are not new, secondary reagent cross-reactivity can happen when multiple sequential rounds of IHC are performed. Given the novelty in the sequential nature of the technique, how the secondary reagents may cross-react has not been studied in detail. This could happen when secondary reagents from two different rounds within a stripping cycle recognize each other, leading to a binding strong enough to localize the active peroxidase where the inactivated one from the previous round is. The end result is signal formation from both rounds (Figure 5A). For example, when we started including goat anti-human primary antibodies in our lymph node scIHC panel, we detected them with a rabbit anti-goat secondary HRP polymer (Table S2 and S3). If a rabbit and goat primary antibodies were to be used within the same stripping cycle, then the two secondary HRP polymers would bind to each other, leading to cross-reactivity (Figure 5B). Depending on the markers involved, this issue can be extremely difficult to realize and often remains unnoticed, leading to confounding results and even when noticed, it limits the choice of species that can be used within a stripping cycle (in the example above, either rabbit or goat), leading to bigger complications in defining the biomarker order for the panel. To avoid this, whenever a validated primary antibody made in goat is added to a cycle, a horse anti-rabbit secondary reagent should be used in that same cycle in place of the goat anti-rabbit secondary HRP polymer. This avoids cross-reactivity between secondary reagents (Figure S1). Of note, cross-reactivity does not happen when only one of the two secondary reagents can recognize the other: for example, a rabbit anti-goat and a goat anti-rat secondary HRP polymers (Figure 5C). We have summarized these cross-reactivity concerns and additional cost-effective analysis in a “decision table” to determine antibody order within a stripping cycle (Table S3 and S4).

**Figure 5.**
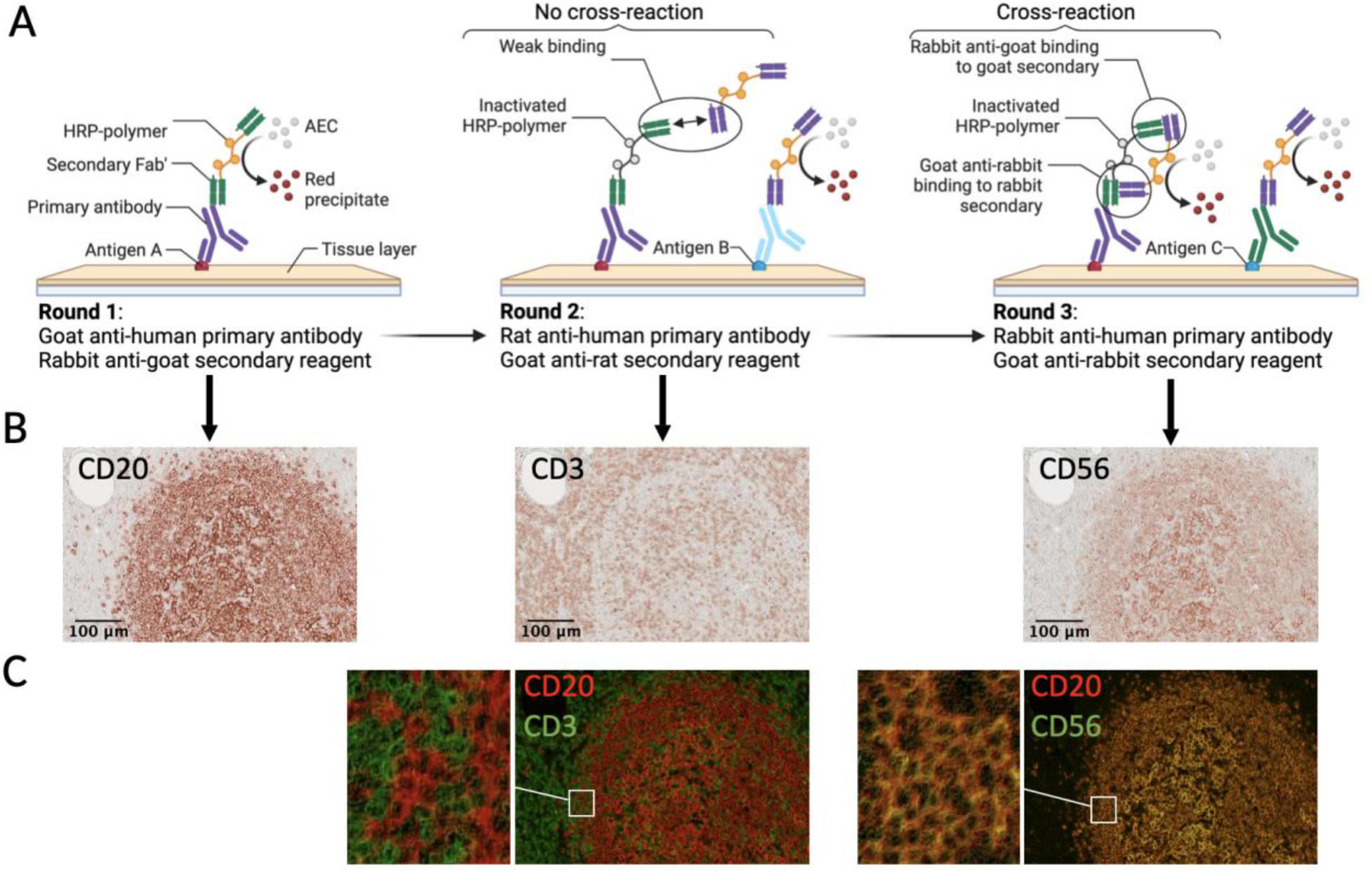
Appropriate selection of secondary antibodies is critical to avoid cross-reactivity. Schematic of a cross-reaction (A) and practical examples (B-C). Using a rabbit anti-goat secondary reagent in the first round of a cycle will result in cross reactivity with a secondary reagent that is goat anti-rabbit (round 3) likely because of cross-binding between the reagents. If only one of the secondary reagents can bind the other, no cross-reactivity is observed (middle panels). Pseudo-colored overlays (C) show cross-reaction (right panel) or absence thereof (left panel). Scale bar: 100 um (10X).

### 3.5 Antibody stripping versus antigen retrieval

Antibody stripping is the process of removing bound primary antibodies from the tissue. It is a necessary step in scIHC, not only because it allows to assess additional biomarkers, but also because the treatment to strip primary antibodies also unmasks antigenic epitopes for better detection. However, it is important to distinguish the two procedures since they have different purposes. Importantly, selecting the appropriate antigen retrieval technique depends on a variety of variables such as the method and duration of fixation, the type of tissue, the target antigen and the antibody used [33]. Most of the antibodies we selected performed optimally using a citrate-based antigen retrieval buffer, pH 6.0. However, we had one antibody, BCL6, which did not perform well in acidic antigen retrieval conditions. Instead, this antibody performed optimally in a Tris-EDTA antigen retrieval buffer, pH 9.0. To assess if this buffer can efficiently strip primary antibodies from previous cycle (which include CD68), we imaged the section after AEC treatment (without re-staining with primary/secondary antibodies). We observed carryover from the previous cycle (Figure 6A-B). In order to properly strip primary antibodies from previous cycle and unmask BCL6 epitopes, we stripped the slides with citrate buffer, allowed to cool to room temperature, and then performed antigen retrieval with Tris-EDTA buffer. This procedure was necessary to detect the appropriate signal of BCL6 without the carryover of CD68 from the previous cycle (Figure 6C). The complete absence of carryover between sequential rounds of staining is demonstrated by: i) CD68+ cells from panel A can be easily distinguished from BCL6+ cells found in their vicinity, in panel C, because their morphologies are very different; ii) CD68+ cells are not stained in panel C; iii) BCL6+ cells clustered at the center of the follicle are stained in panel C but not in panel A; iv) EDTA-based antigen retrieval did not fully strip anti-CD68 antibodies, which can still be detected in panel B, after incubation with AEC. Of note, biomarkers that require basic-pH antigen retrieval can be placed at the beginning of the panel, which avoids epitopes on these biomarkers to undergo unnecessary stripping cycles.

**Figure 6.**
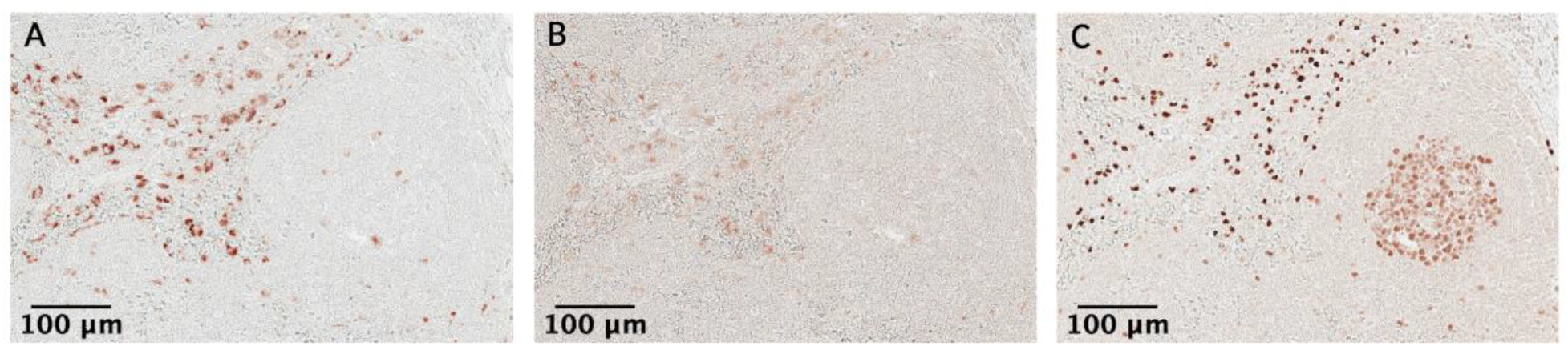
Citrate stripping is necessary before EDTA-based antigen retrieval. A patient sample was stained with CD68 in cycle 11 round 2 (A). In the next cycle, an EDTA antigen retrieval buffer was used to detect BCL6, without citrate stripping. Carry over from the previous cycle was observed by treatment with AEC and visualization (B). Stripping with a citrate antigen retrieval buffer was performed and EDTA-based antigen retrieval was conducted again, which allowed to visualize BCL6 without carry over from previous staining (C). Note the different morphology of BCL6+ and CD68+ cells within the same region (A, C). Same section is shown in (A-C). Scale bar: 100 um (10X).

### 3.6 Spatial analysis of lymph node cells

Digitalization of image data promises to accelerate our understanding of complex biological systems [9, 34–37]. However, each imaging technique presents unique challenges in the quantification of image data. To extract the highest amount of digital data without sacrificing data quality and accessibility, we partnered with Ariadne, a company specialized in advanced image analysis. The main steps of the analysis pipeline we developed are: alignment, segmentation, location and intensity extraction, subset identification and spatial analysis (Figure 7A). As an example of the type of information that can be extracted by digitalizing histological images, we focused on antibody producing cells known as plasmablasts. To identify potentially meaningful interactions with other immune subsets, we calculated the distance between each plasmablast and cells expressing antibody receptors (that is, Fc receptors). These cells are mostly of myeloid origin, with the exception of NK cells [38]. We observed that plasmacytoid dendritic cells (pDCs) were often in contact with plasmablasts, as evidenced by the fact that almost 50% of them was within 10 um (center to center) from pDCs (Figure 7B). When we visually inspected these two immune populations, we observed that they are often found closely interacting with each other (Figure 7C). By comparing the locations of plasmablasts with respect to pDCs at a tissue scale, we found that indeed they have contact interactions in both the subcapsular and medullary regions (Figure 7D). This was not the case for sinusoidal macrophages or granulocytes. Importantly, the frequence of contact interactions was not dependent on cell abundance (Figure 7D). Overall, the quantitative analysis we developed allows to study the cellular composition and location of the tissue under investigation.

**Figure 7.**
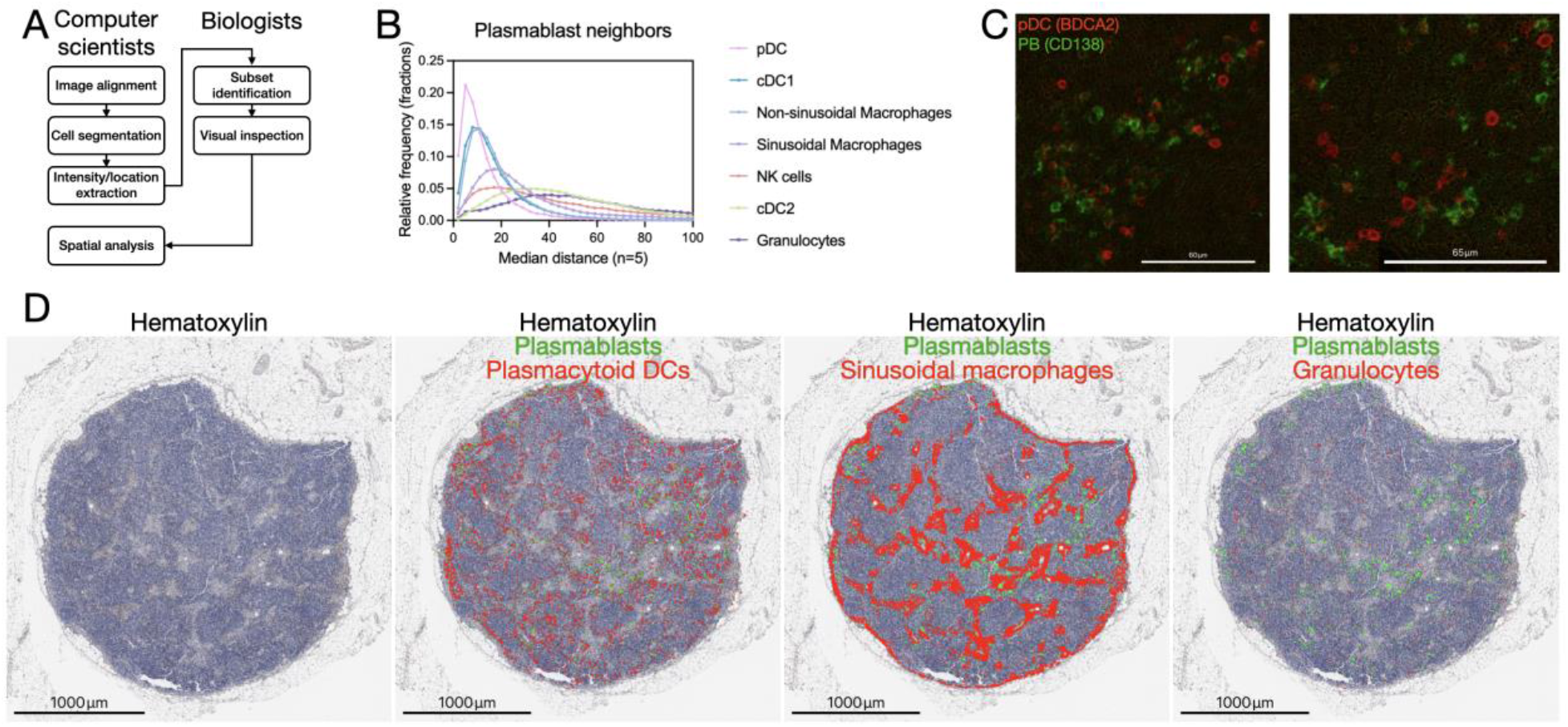
Analysis of plasmablast neighborhood. Schematic of the main steps of the analysis pipeline (A). Frequency distribution of the distance between plasmablasts and the indicated immune cell types (B). Micrographs showing two representative fields rich in plasmablast–plasmacytoid dendritic cell contacts (C). Low-magnification view of hematoxylin-stained lymph node superimposed with locations of plasmablasts (green dots) and either (D): plasmacytoid dendritic cells, sinusoidal macrophages and granulocytes (red dots). Scale bar: 1000um (10x).

## 4. Discussion

In this work, we provide key information to users aiming to adopt sequential chromogenic IHC as the approach of choice for spatial analyses. We describe how to order biomarkers within a cycle, how to carefully choose secondary HRP polymers and we discuss which situations recommend doubling of the endogenous peroxidase blocking step. Overall, these considerations effectively increased biomarker detection accuracy while eliminating issues like carryover and cross reactivity.

Sequential chromogenic IHC has several advantages over fluorescence-based techniques: it does not require balancing fluorochrome brightness with marker abundance [39], provides higher sensitivity of detection, avoiding the need for signal amplification [40], up to 15-20 patient samples can be processed by an operator at a time, pathology departments are already equipped to perform scIHC (including image scanners), and generally lower entry costs. Despite its advantages, scIHC is not without caveats. This technique can be relatively time consuming because only one or two biomarkers can be assessed at the same time (eg. by using a red and blue chromogen). Obtaining data from one staining round can take 4- 24 hours to generate, depending on incubation times. Thus, the maximum daily output is 4 biomarkers (that is, two staining rounds per day). As we consider in this manuscript, there are constraints in arranging antibodies within a panel, so that antibodies that are sensitive to repeated stripping cycles need to be employed first. While these caveats should not discourage use of this technique, they should be taken into account when determining feasibility and design of experiments. As it is the case for other sequential histological approaches, operators must pay special attention to the expected distribution of signal for current biomarker and compare it to previous biomarkers in order to detect potential issues with panel design. Expected signal distribution for the tissue under investigation can be learnt from Protein Atlas website (https://www.proteinatlas.org/).

The procedure we describe can be upgraded relatively easily by multiplexing two chromogens, which can further increase the throughput without sacrificing quality. To multiplex the scIHC with two chromogens, two different primary antibodies (from different species) can be employed within each round, followed by the corresponding secondary detection reagents conjugated to different enzymes (for example, horseradish peroxidase and alkaline phosphatase). These two enzymes will generate precipitate of different color (red and blue, in the example above), that can be imaged at the same time. Simple computational approaches would then separate the signals from the two biomarkers, effectively doubling the throughput of the technique.

A major hurdle in any sequential histological approach is the analysis pipeline. We decided to outsource the development of the analysis pipeline to professionals with established experience in the field [41]. By closely collaborating with computational biologists at Ariadne.ai (https://ariadne.ai), we developed a pipeline based on commonly used steps [11] but implementing cutting-edge computational approaches, including non-rigid tissue deformation correction and cellpose-inspired automated cell segmentation (see methods). The result is a plug-and-play approach able to truthfully identify the localization of different cell subsets, which is a pre-requisite for spatial studies. A key factor in the success of the pipeline development was frequent iteration between wet and dry lab scientists, which allowed to quickly spot issues and identify solutions to correct them. For example, since biomarkers by definition work only in the tissue where they have been defined, having a mix of tissues on the slide (for example, lymph node, fat and connective tissues) leads to the inclusion of spurious cells within certain biomarkers (eg. CD141, which labels both cDC1 and endothelial cells). To fix this issue, we defined a tissue of interest mask that excludes segmented cells not pertaining to the lymph node tissue proper. By exporting the biomarker intensities and cell locations in FCS format, we were able to avoid the use of FCSExpress (Denovo software), currently the only commercial software able to handle digitalized image data. We replaced the live image gating inspection in FCSExpress by importing gated events into KNOSSOS, which runs without crashing and is not affected by frequent software bugs. KNOSSOS is developed by the Max Planck Institute in Germany, and it can display tissue masks, segmented cells and chromogenic signals (including hematoxylin) all together or individually, as layers. Although the neural network-based cell segmentation of a densely cellularized tissue like the lymph node worked very well in distinguishing both round and elongated nuclei (Movie S1), a future improvement of our pipeline will be to consider chromogenic signals during segmentation, in order to avoid spillover of said signals into neighboring cells, which would be especially important for membrane biomarkers.

As proof of concept of what can be discovered by implementing our procedures and bio-informatic pipeline, we present novel data on the neighborhood of lymph node resident plasmablasts. To our surprise, we identified plasmacytoid dendritic cells (pDCs) as the cell type with contact interactions with plasmablasts in head and neck cancer patient lymph nodes. The communication between these two immune subsets has been reported only *in vitro* [42, 43], and here we confirm that plasmablasts and pDCs display contact interactions *in vivo.* Given the role of TLR receptors (highly expressed in plasmacytoid dendritic cells) in vaccine efficacy [42, 44–46], the significance of such interaction decisively warrants further studies.

## 5. Supplementary Material

**Figure S1.**
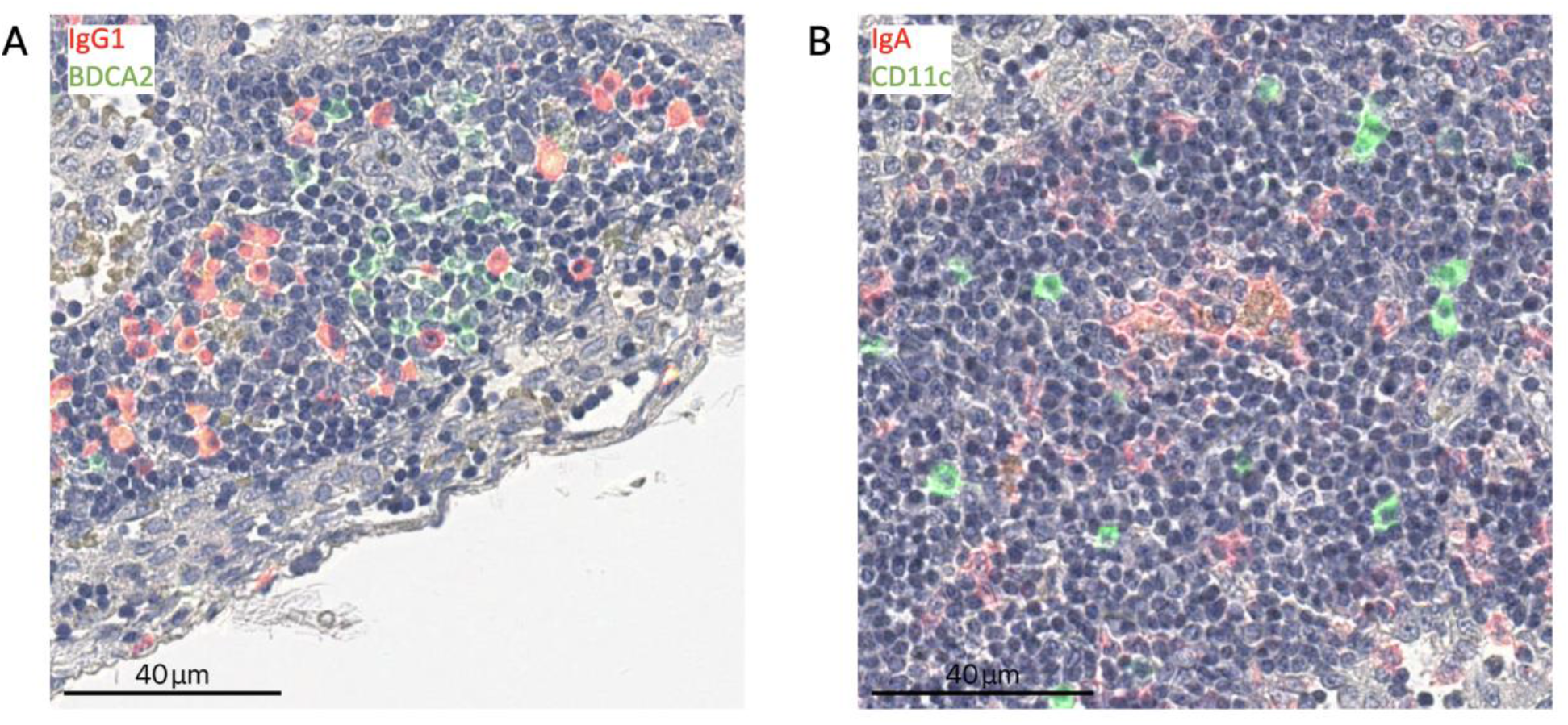
Avoid cross-reactivity of secondary reagents. Using a horse anti-rabbit (see Figure 3) to detect IgG1 (A) or CD11c (B) does not lead to cross reactivity with the rabbit anti-goat secondary reagent used in the next round to detect BDCA2 and IgA, respectively.

**Figure S2.**
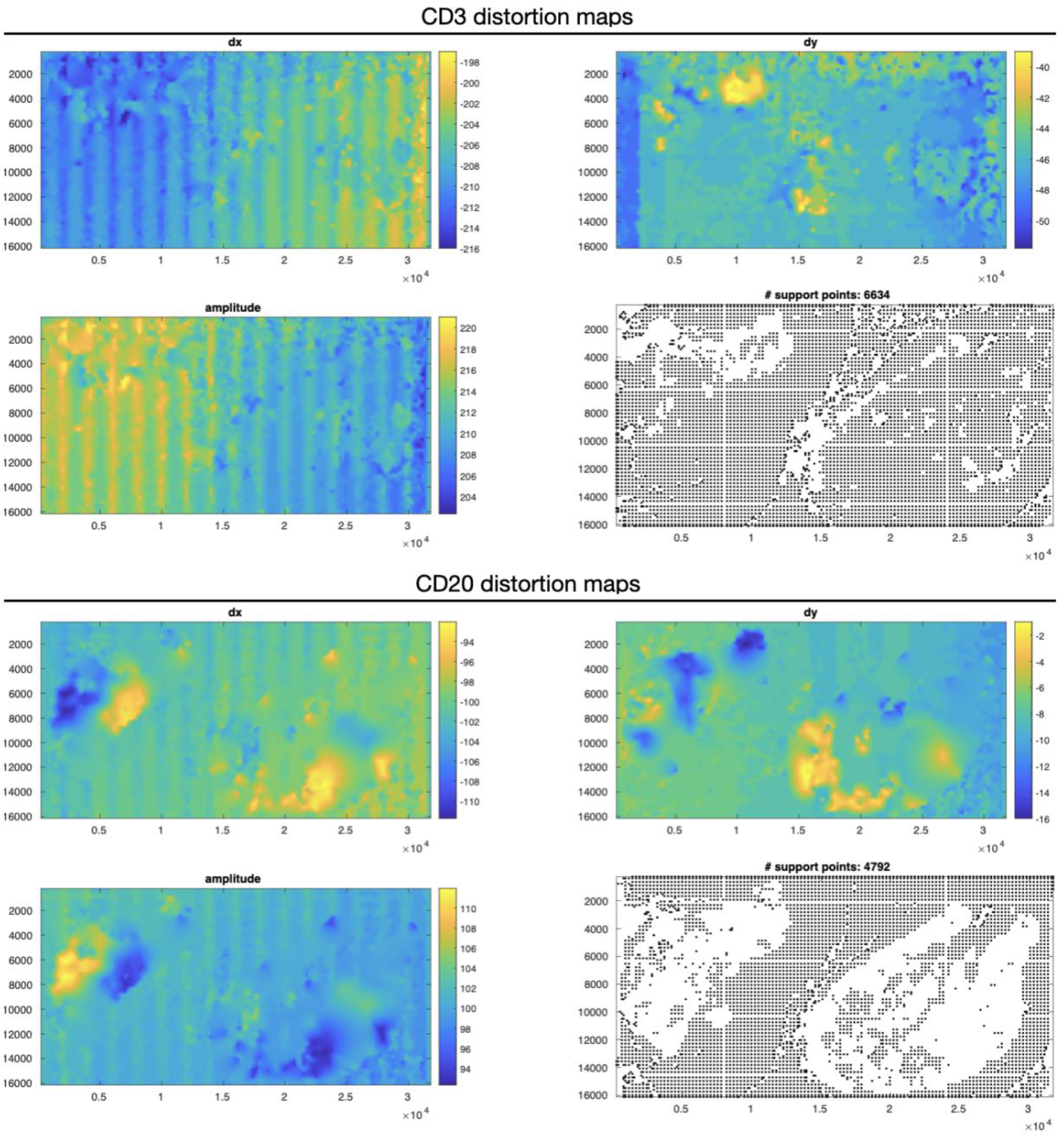
Distortion maps. Two examples of the calculated shifts (dx, dy and amplitude) and support points for CD3 and CD20.

**Figure S3.**
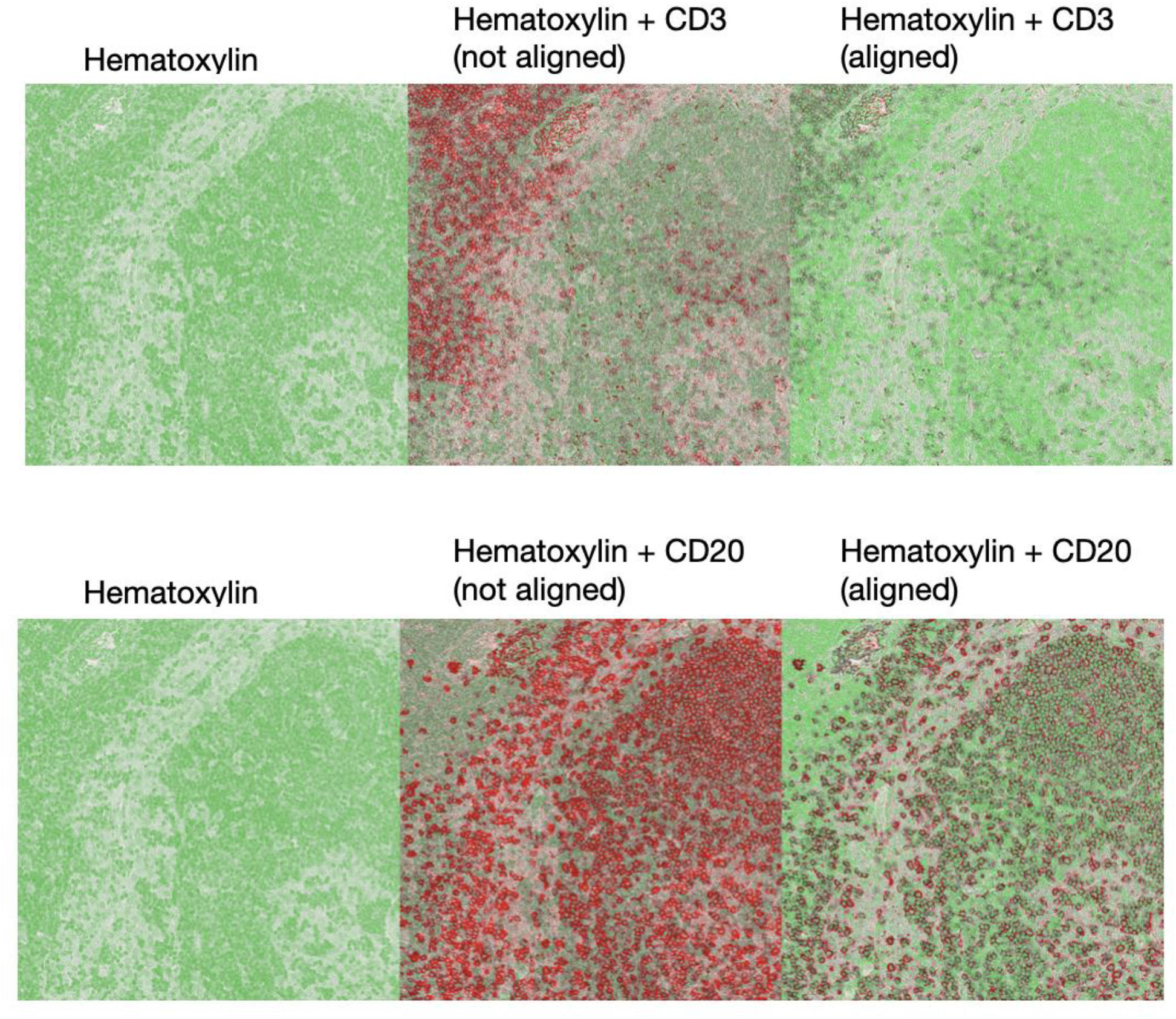
Alignment results. Comparison of before and after alignment for CD3 and CD20 (red) superimposed with hematoxylin (green).

**Figure S4.**
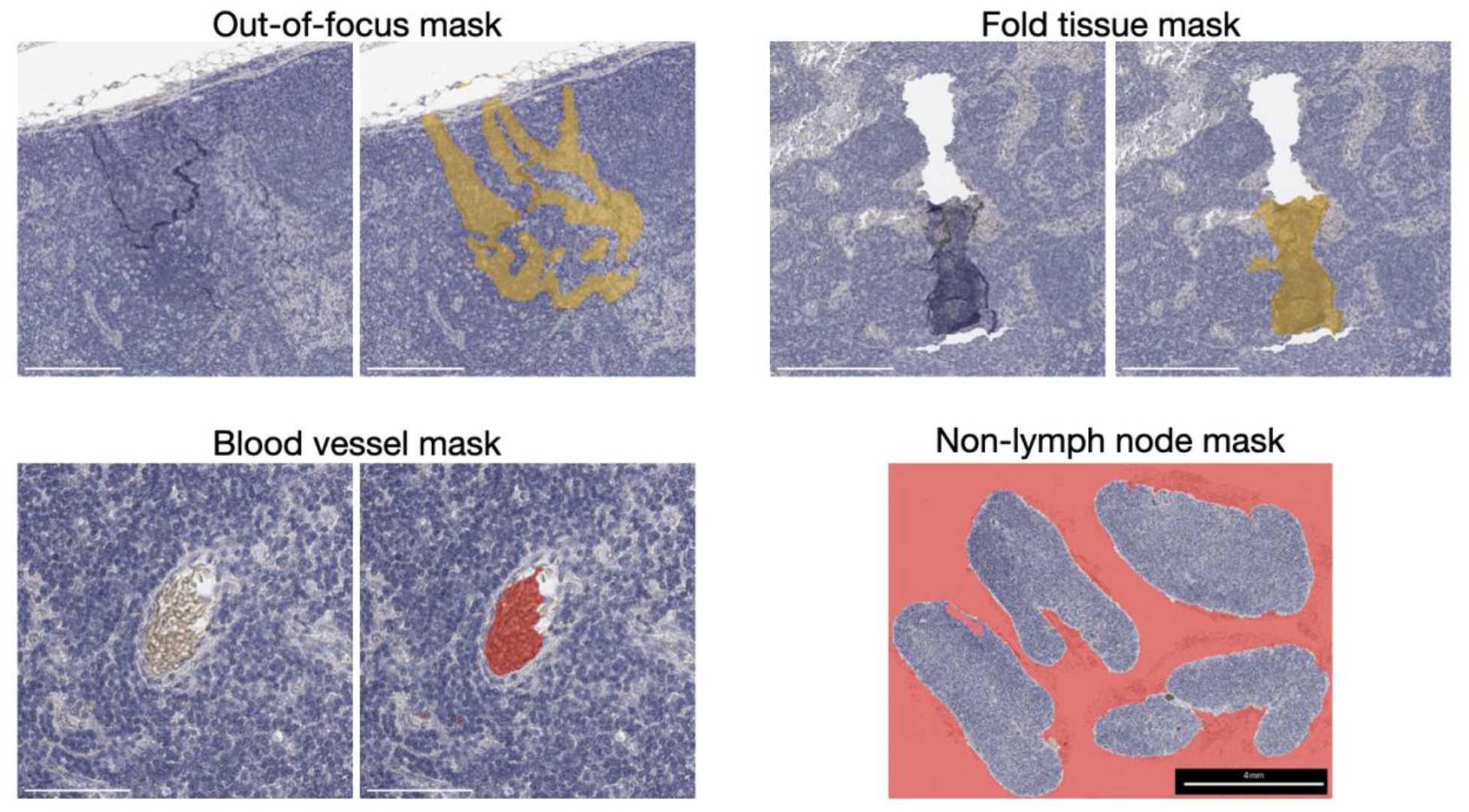
Trained networks detected artifacts and tissue of interest limits, and excluded them from the analysis.

**Table S1.** Sequential IHC panel information.

| Marker | Host Species | Dilution | Reaction | Clone | Vendor | Cell type |
| --- | --- | --- | --- | --- | --- | --- |
| CD56 | Rabbit IgG | 1:500 | Overnight<br>4°C | MA1-06801 | Novus Biologicals | NK cells |
| CD20 | Goat IgG | 1:500 | Room temp,<br>30 min | polyclonal | Abcam | B cells |
| CD66b | Mouse IgG | 1:600 | Room temp,<br>30 min | G10F5 | BD Biosciences | Granulocytes |
| PD1 | Goat IgG | 1:200 | Overnight<br>4°C | polyclonal | R&D Systems | T follicular<br>helper |
| IgM | Mouse IgG | 1:200 | Overnight<br>4°C | IM260 | Novus<br>Biologicals | Multiple |
| CD3 | Rat IgG | 1:250 | Room temp,<br>30 min | CD3-12 | Abcam | T cells |
| IgE | Goat IgG | 1:1000 | Room temp,<br>30 min | polyclonal | Novus<br>Biologicals | Multiple |
| HLA-DR/DP/DQ | Mouse IgG | 1:500 | Room temp,<br>30 min | WR18 | LSBio | Multiple |
| CD169 | Sheep IgG | 1:200 | Room temp,<br>30 min | polyclonal | R&D Systems | Sinusoidal<br>macrophages |
| CD11c | Rabbit IgG | 1:200 | Room temp,<br>30 min | SI19-06 | Novus<br>Biologicals | Dendritic cells |
| IgA | Goat IgG | 1:10,000 | Overnight<br>4°C | polyclonal | Novus<br>Biologicals | Multiple |
| IgG1 | Rabbit IgG | 1:500 | Room temp,<br>30 min | RM117 | Novus<br>Biologicals | Multiple |
| BDCA2 | Goat IgG | 1:100 | Room temp,<br>1 hr | polyclonal | Novus<br>Biologicals | Plasmacytoid<br>dendritic cells |
| CD138 | Mouse IgG | 1:50 | Room temp,<br>30 min | MI15 | ThermoFisher | Plasmablasts<br>and plasmacells |
| CD38 | Mouse IgG | 1:20 | Room temp,<br>1 hr | 38C03<br>(SP32) | ThermoFisher | Plasmacells |
| CD88 | Rabbit IgG | 1:500 | Overnight<br>4°C | polyclonal | Novus<br>Biologicals | Recirculating<br>monocytes |
| CD141 | Sheep IgG | 1:20 | Room temp,<br>2 hr | polyclonal | Novus<br>Biologicals | Type-1<br>conventional<br>DCs |
| CD1c | Mouse IgG | 1:200 | Room temp,<br>1 hr | OTI2F4 | Novus<br>Biologicals | Type-2<br>conventional<br>DCs |
| IgG3 | Rabbit IgG | 1:500 | Overnight<br>4°C | RM119 | Novus<br>Biologicals | Multiple |
| PanCK | Rabbit IgG | 1:500 | Room temp,<br>1 hr | polyclonal | Novus<br>Biologicals | Squamous<br>carcinoma cells |
| CD4 | Mouse IgG | 1:150 | Room temp,<br>30 min | OT12H8 | LSBio | CD4 T cells |
| FDC | Mouse IgG | 1:1000 | Overnight<br>4°C | CNA.42 | Novus<br>Biologicals | Follicular<br>dendritic cells |
| Ki67 | Rabbit IgG | 1:500 | Room temp,<br>30 min | SP6 | MilliporeSigma | Proliferation |
| CD8 | Mouse IgG | 1:50 | Room temp,<br>30 min | C8/144B | ThermoFisher | CD8 T cells |
| Tbet | Rabbit IgG | 1:200 | Room temp,<br>1 hr | D6N8B | Cell Signaling | Activation |
| CD68 | Mouse IgG | 1:2000 | Room temp,<br>30 min | KP1 | Novus<br>Biologicals | Macrophages |
| BCL6 | Rabbit IgG | 1:50 | Room temp,<br>1 hr | polyclonal | Novus<br>Biologicals | Multiple |

**Table S2.** Secondary HRP Round Considerations.

| Primary Antibody Species | Secondary HRP Polymer | Manufacturer | Round Considerations |
| --- | --- | --- | --- |
| Rabbit | Horse αRabbit | Vector Labs | For cycles that have goat derived antibodies |
| Rabbit | Goat αRabbit | Nacalai USA | For cycles that do not have goat derived antibodies* |
| Goat | Rabbit αGoat | Nacalai USA | Place goat antibody before mouse and rat derived antibodies |
| Mouse | Goat αMouse | Nacalai USA | Use in any round, but after goat derived antibodies |
| Rat | Goat αRat | Nacalai USA | Use in any round, but after goat derived antibodies |
| Sheep | Donkey αSheep | R&D Systems | Use in any round |

\*Due to cost consideration of Horse $\alpha$ Rabbit HRP polymer at the time of experimentation.

**Table S3.** Comprehensive database information for mIHC panel development.

| Database Column | Information captured |
| --- | --- |
| Marker | What the antibody detects |
| Species | Species the antibody was produced in: Mouse, Rabbit, Rat, Goat, Sheep |
| Reactivity | Species the antibody reacts to: Human |
| Dilution | Specific dilution the antibody was tested |
| Incubation | 30 minutes – 2 hours a room temperature, overnight at 4°C |
| AEC incubation | The amount of time the slides was incubated with visualization buffer (this was used to inform planning and timing of protocol) |
| Protein Block incubation | Amount of time protein block was incubated (at room temperature) |
| Antigen Retrieval | Citrate (pH 6.0) or Tris-EDTA (pH 9.0) |
| Slide ID | Unique identifier of slide |
| Cycle | The specific cycle number the slide was tested: C01, C02, etc. |
| Number of Rounds | The cumulative number of rounds a unique slide has been subjected to irrespective of cycle. |
| Worked | Yes, No, Yes with background, Yes with faint staining, Background only, No with carryover |
| Tissue Type | Specific type of tissue: Lymph node, tonsil, skin |
| Experiment | The name of the experiment and data location (to access images) |
| Clone | Specific clone of antibody |
| Company | Manufacturer of antibody |
| Catalog Number | Catalog number |

**Table S4.** Bullet list of considerations to develop a biomarker panel.

|  |  |
| --- | --- |
| 1 | Define a wish list of biomarkers |
| 2 | Identify potential alternative biomarkers (in case some primary antibodies need to be replaced, see step 5 below) |
| 3 | Consult our antibody database (and other sources) to determine the staining order by collecting information on which cycle the selected primary antibodies will work in and start filling the panel |
| 4 | Identify conflicts (eg. 2 or more same-species primary antibodies need to be in the same cycle) and unknowns (that is, primary antibodies without information on staining quality by cycle) |
| 5 | Find alternative primary antibodies to resolve the conflicts (including using alternative biomarkers) |
| 6 | Test newly-adopted (unknowns) primary antibodies on practice slide to identify saturating dilution, pH of antigen retrieval step and latest cycle in which performance is maintained (or the target cycle within a tentative panel) |
| 7 | Run the final panel on test samples and assess potential cross-reactivity, carry over by comparison of signals within a stripping cycle |

Movie S1. Cell segmentation of hematoxylin-stained tissue. Example of segmented cells in the sub- capsular region, showing both small round nuclei in lymphocyte-rich areas (B cell follicle, top left) and elongated nuclei in macrophage-rich areas (sub-capsular sinus, middle).

